# PyRanges: efficient comparison of genomic intervals in Python

**DOI:** 10.1101/609396

**Authors:** Endre Bakken Stovner, Pål Sætrom

## Abstract

**Summary:** Complex genomic analyses often use sequences of simple set operations like intersection, overlap, and nearest on genomic intervals. These operations, coupled with some custom programming, allow a wide range of analyses to be performed. To this end, we have written PyRanges, a data structure for representing and manipulating genomic intervals and their associated data in Python. Run single-threaded on binary set operations, PyRanges is in median 2.3-9.6 times faster than the popular R GenomicRanges library and is equally memory efficient; run multi-threaded on 8 cores, our library is up to 123 times faster. PyRanges is therefore ideally suited both for individual analyses and as a foundation for future genomic libraries in Python.

**Availability:** PyRanges is available open-source under the MIT license at https://github.com/biocore-NTNU/pyranges and documentation exists at https://biocore-NTNU.github.io/pyranges/

**Supplementary information:** Supplementary data are available.

## 1. Introduction

Comparing sets of intervals is a fundamental task in genomics, and a few basic operations allow for answering many scientific questions. For example, to find genes potentially targeted by a transcription factor, one can intersect the sets of intervals representing gene positions and representing transcription factor binding sites to identify those that overlap.

Several toolboxes of genomic operations exist, such as bedtools (Quinlan et al., 2010) and bedops (Neph et al., 2012) for the command line and GenomicRanges (Lawrence et al., 2004) for the R programming environment.

GenomicRanges is a data structure for representing and operating on genomic intervals and their metadata, which are stored as a 2D-table in memory. By providing methods for access and for set operations on genomic intervals, programmers can use the R programming language to manipulate and analyse the contents of GenomicRanges. Consequently, GenomicRanges is a powerful tool for writing complex and custom genome analyses. Indeed, in R, GenomicRanges is a foundational library, and a cornerstone of genomics packages in the R Bioconductor project (Gentleman, 2004).

Python is currently ranked as the most popular programming language in the world (according to IEEE Spectrum’s compound metric; S.Cass et. al, 2018) and is much used in data science and bioinformatics, yet it lacks a GenomicRanges implementation. The PyRanges library remedies this by providing a version which is multithreaded, fast, and memory-efficient.

## 2. Library

### 1. Implementation

The PyRanges data structure is logically represented as a 2D-table. Each row represents an interval, and the columns each describe either a part of the location (chromosome, start position, end position, and optionally, strand) or metadata (name, score, exon number or any arbitrary value desired by the user). The underlying implementation uses a dictionary that maps chromosome and strand pairs to their respective 2D-tables; however, this division is largely invisible to the user. The data in the 2D-tables are stored in Pandas DataFrames, thus allowing the vast Python science stack to be used seamlessly with PyRanges. Furthermore, DataFrames allow for storing the data contiguously in native data types, such as integers, floats, or categoricals, to ensure memory-efficiency.

To make PyRanges fast, its operations are written in Cython or C. Moreover, by keeping the data belonging to each chromosome in separate DataFrames, these logically distinct data can easily be independently processed. For single threaded processing, this implementation detail has limited effect, but for multi threaded processing, we avoid the substantial time costs of splitting and merging the data for each operation. PyRanges provides parallel processing through the Ray framework (P.Moritz et al., 2017), resulting in a speedup provided the data are sufficiently big (see timings).

### 2. Functionality

PyRanges main functionality includes functions for reading genomic intervals from files, and unary and binary functions for manipulating one and two sets of genomic intervals. File reading functions support common formats such as bed, GTF/GFF, and bam. Unary functions manipulate single PyRanges by subsetting, clustering, or computing coverage; that is, the number of intervals overlapping each genomic position. Binary functions include operations such as intersection, nearest, and subtract that create a new set of genomic intervals by comparing two sets of intervals. See the Supplementary text for a full list of PyRanges’ operations.

PyRanges also uses and provides two stand-alone libraries useful beyond bioinformatics. One library (pyrle) implements run-length encoding arithmetic, which is useful to compactly represent and efficiently do arithmetic on the coverage (or any other nucleotide-associated score) of sets of regions. The other library (NCLS) implements the Nested Containment List, which is an immutable interval-tree with better memory-efficiency and speed than a regular interval-tree both for tree construction and interval queries (see Supplementary Timings).

### 3. Performance

The PyRanges library has been extensively benchmarked for both speed and memory use (Fig 1; Supplementary Timings). We used two types of data for testing: 1) libraries of reads only, i.e. they included no metadata and were hence more lightweight and 2) GTF annotations. We used unsorted test files generated by bedtools random for hg38 to simulate the read files. To create a large GTF we used sampling with replacement on the Gencode hg38 GTF.

**Fig. 1.**
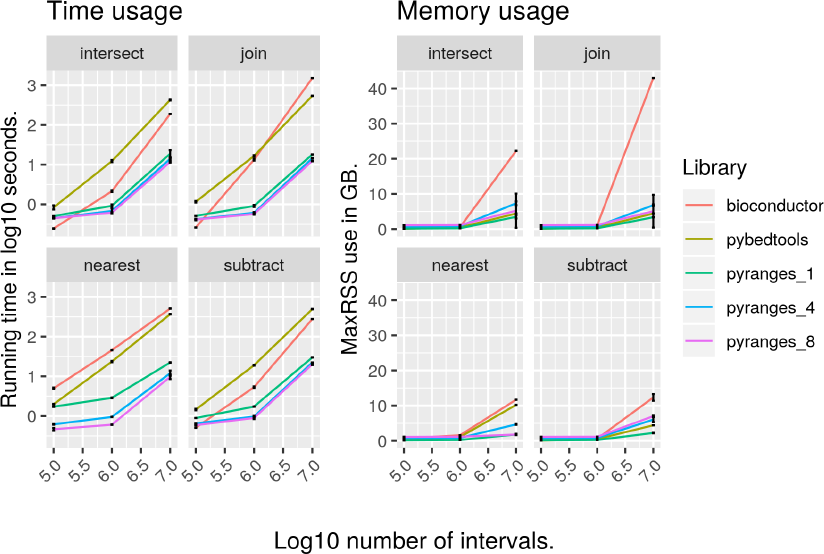
(Left) Running time and (Right) memory usage as a function of the number of intervals for four common binary functions on genomic intervals; see Supplementary Timings for complete benchmark results.

For binary operations, PyRanges in single-threaded mode was 6.5 - 31 (median 14) and 9.8 - 36 (median 24) times faster than pybedtools on 1e6 and 1e7 intervals, respectively. Compared to GenomcRanges, PyRanges was 1.3 - 16 (median 2.3) and 1.9 - 84 (median 9.6) times faster. Run multi-threaded on 8 cores, the speed-ups for the same operations on 1e7 intervals were 13 - 63 and 1.8 - 123 times compared to pybedtools and GenomicRanges, respectively. For all operations, PyRanges run single-threaded on 1e7 intervals had a median speed-up of 26 and 4.0 times and used a median 5.7 and 5.1 times less memory compared to pybedtools and GenomicRanges, respectively.

## 3. Conclusion

PyRanges is an efficient and feature rich library for genomics in the extremely popular Python programming language, and the only one of its kind. We therefore expect it to be a boon to current and future bioinformaticians and researchers working in Python.

## Supporting information

Supplementaries: discussion

Supplementaries: time and memory use

