## Supplementaries: discussion for "PyRanges: efficient comparison of genomic intervals in Python"

### PyRanges supplementary text 2

This supplementary text includes additional comparisons between PyRanges and the pybedtools and GenomicRanges libraries, descriptions of general-purpose stand-alone libraries developed to support PyRanges, and descriptions of the PyRanges testing and continuous integration frameworks.

#### PyRanges vs pybedtools

##### PyRanges is all Python, pybedtools is not

The PyRanges package is the first Python-native implementation of a general-purpose, high-performance genomics library. Its nearest competitor, pybedtools merely wraps the command line tool BEDTools and makes its functions available from Python. As such, pybedtools relies on reading and writing temporary files on disk, which makes it slow, prone to filesystem-related errors, and reliant upon adequate disk storage.

##### PyRanges is all in-memory, pybedtools uses disk

As pybedtools stores data in files instead of memory, combining pybedtools with other data processing tools is cumbersome and inefficient. For example, to use the Python data science stack on the data in combination with pybedtools, one needs to read the pybedtools data into memory, operate on the data, and then write the resulting data back to disk. The alternative way is to operate on the data while pybedtools processes it, but this approach is very slow. The reason is that the approach requires one to read each line individually from pybedtools, turn the line into Python data, operate on the data, and then convert the result into a string and write it to an output file. PyRanges is already orders of magnitude faster than pybedtools; doing such custom analyses on the data would greatly increase the difference.

##### pybedtools is strict about the file format, PyRanges is not

Lastly, pybedtools is strict about the data format. It can only read and use gtf, bam, and bed files. Furthermore, its strict adherence to formats means that if for example a tab-delimited file contains strand info, but not in the sixth column, stranded operations will silently fail without a warning.

```
head f1.bed f2.bed
==> f1.bed <==
chr1    1   2   a   0   +
```

```
==> f2.bed <==
chr1    1   2   a   0   ReadName    -
bedtools intersect -S -a f1.bed -b f2.bed
# no output
```

This is unfortunate, as bioinformatics has an abundance of file formats for different software packages. PyRanges is not strict about formats at all; as long as the data has columns for Chromosomes, Starts and Ends it will work. If a stranded operation is requested, but no strand is present it will fail with an explicit error.

##### PyRanges are easy to query, pybedtool-objects are not

Another disadvantage of pybedtools is that querying the data for interactive or exploratory analysis is hard. If you for example want to know all intervals on chromosome 6 between nucleotides x and y on the reverse strand, you need to create a bed file with the info (including dummy columns for name and score) and then run an intersect operation with your original file and the single interval file made explicitly for a query. This is inordinately time-consuming and inconvenient.

In comparison, PyRanges uses Python's getitem syntax to allow you simply to inspect the data in position-based ways:

```
import pyranges as pr

gr = pr.db.ucsc.genes("hg38") # fetch using mysql

gr["chr6"]
# +--------------|-----------|-----------|----------|------------|----------------|---------
# | Chromosome   | Start     | End       | Strand   | Feature    | TranscriptID   | ExonNum
# | (int16)      | (int32)   | (int32)   | (int8)   | (int8)     | (int32)        | (int16)
# |--------------|-----------|-----------|----------|------------|----------------|---------
# | chr6         | 292056    | 351355    | +        | transcript | NM_001286555   | nan
# | chr6         | 292056    | 351355    | +        | transcript | NM_020185      | nan
# | chr6         | 292056    | 351355    | +        | transcript | NR_104473      | nan
# | ...          | ...       | ...       | ...      | ...        | ...            | ...
# | chr6         | 233200153 | 233200264 | -        | exon       | NM_144781      | 1
# | chr6         | 233200153 | 233200264 | -        | exon       | NM_144781      | 2
# | chr6         | 233200153 | 233200264 | -        | exon       | NM_144781      | 3
# +--------------|-----------|-----------|----------|------------|----------------|---------
# PyRanges object has 38507 sequences from 1 chromosomes.

gr["chr6", 100000:100000]
# Empty PyRanges
```

```
gr["chr6", 100000:200000]
# +--------------|-----------|-----------|----------|------------|----------------|---------
# | Chromosome   |     Start |       End | Strand   | Feature    | TranscriptID   |   ExonN
# | (int16)      |   (int32) |   (int32) | (int8)   | (int8)     | (int32)        |       (i
# |--------------|-----------|-----------|----------|------------|----------------|---------
# | chr6         |    140263 |    148159 | -        | transcript | NR_109817      |
# | chr6         |    181465 |    205484 | -        | transcript | NR_126020      |
# +--------------|-----------|-----------|----------|------------|----------------|---------
# PyRanges object has 2 sequences from 1 chromosomes.

gr["+"]
# +--------------|-----------|-----------|----------|------------|----------------|---------
# | Chromosome   | Start     | End       | Strand   | Feature    | TranscriptID   | ExonNum
# | (int16)      | (int32)   | (int32)   | (int8)   | (int8)     | (int32)        | (int16)
# |--------------|-----------|-----------|----------|------------|----------------|---------
# | chr1         | 11873     | 12227     | +        | exon       | NR_046018      | 1
# | chr1         | 11873     | 12227     | +        | exon       | NR_046018      | 2
# | chr1         | 11873     | 12227     | +        | exon       | NR_046018      | 3
# | ...          | ...       | ...       | ...      | ...        | ...            | ...
# | chrY         | 57067864  | 57130289  | +        | transcript | NM_005638      | nan
# | chrY         | 57184100  | 57197337  | +        | transcript | NM_002186      | nan
# | chrY         | 57184100  | 57197337  | +        | transcript | NM_176786      | nan
# +--------------|-----------|-----------|----------|------------|----------------|---------
# PyRanges object has 437019 sequences from 330 chromosomes.
```

#### PyRanges vs. the R BioConductor GenomicRanges

PyRanges has many advantages over its cousin in R too. When it comes to purely objective measures, PyRanges is faster, more memory-efficient, and supports using multiple cores. In addition, PyRanges has several advantages from a design and usability standpoint, as discussed in the following sections.

##### All PyRanges methods are contained within the pyranges package or the objects themselves

The R GenomicRanges package clutters the namespace with method-names upon import, whereas all PyRanges methods are either available as methods in the pyranges library (`pr.db.ucsc.genes("hg19")`) or on the objects themselves (`pyrange1.intersect(pyrange2)`).

##### PyRanges allow for method chaining, GenomicRanges do not

Doing consecutive operations on PyRanges objects is easy; for example, given three PyRanges a, b, and c, finding the intervals in c nearest to b and intersecting these with a, can be done as follows:

```
c.nearest(b).intersect(a)
```

In R, these operations require the following code (see the next section for an explanation of why the call to `subjectHits` is required):

```
pintersect(a, c[subjectHits(distanceToNearest(c, b))])
```

The more operations are chained, the more difficult the GenomicRanges code is to write and follow, making the code more prone to bugs.

##### GenomicRanges has an anarchic API, PyRanges does not

Many GenomicRanges-methods do not return new GenomicRanges-objects, but rather a hits object which needs to be queried. This is the reason why a call to `subjectHits` is required in the following code:

```
pintersect(a, c[subjectHits(distanceToNearest(c, b))])
```

In comparison, the following code

```
pintersect(a, distanceToNearest(c, b))
```

would lead to the error

```
Error in (function (classes, fdef, mtable)  :
  unable to find an inherited method for function ‘pintersect’ for signature ‘"GRanges", "So
```

##### GenomicRanges-objects are hard to explore

Like pybedtools, GenomicRanges has no method that allows one to query the objects at certain positions. Indeed, the proposed way to do it is to create another object and intersect or use R’s built-in subsetting mechanism based on Boolean expressions: https://www.biostars.org/p/351377/

#### Other useful libraries included in PyRanges

In addition to the PyRanges library, we have written four separate libraries used in PyRanges: sorted_nearest, a library for nearest queries on interval collections; bamread, a library to read data into contiguous, native datatypes; and NCLS and Pyrle, described below. We have made these available as stand-alone libraries because they perform useful functions on their own. ### Nested containment list - NCLS

The NCLS is an optimized implementation of the nested containment list, which is basically an immutable and extremely fast interval tree. We extracted the C-code used from its original implementation in the now defunct PyGR library and wrote a Cython wrapper for it, allowing seamless use from Python. We also implemented several optimizations such as removing interval-level variables not needed for pure overlap-queries, implemented both 32 and 64-bit versions and wrote methods which allow batch-querying entire collections of intervals against each other at native speed. On a test datafile of 100 million intervals the NCLS is built in 3.15 seconds;in comparison, the interval tree library quicksect, which to our knowledge is the fastest Python alternative, uses 168 seconds on the same data. And when querying the data structure for overlaps with another 100 million reads, NCLS uses 7.2 seconds whereas quicksect uses 148 seconds. As such, our heavily optimized NCLS is 20-50 times faster than the currently fastest Python range overlap query tool. Since NCLS is a library that solves a general problem we expect it to be used outside the scope of genomic intervals.

##### Arithmetic run length encoding - pyrle

The Pyrle library implements arithmetic run length encoding for Python, similar to the R library S4Vectors. An RLE is a compact representation of a vector where a lot of consecutive elements have the same value. The coverage of sequencing reads within a chromosome is, for example, often convenient to represent in RLE as such a coverage has long stretches of 0, places without coverage, or many repeating elements. Using such a representation, the pileup of reads or the coverage of a genome, often a vector with a length of billions of nucleotides can be represented very compactly. Moreover, arithmetic operations on RLE representations can be performed extremely quickly.

The pyrle library also implements PyRles, a collection of Rle objects, which allow for easy arithmetic on whole genomes. The following code snippets illustrates its use,

```
import pyranges as pr
gr19 = pr.db.ucsc.genes("hg19")
gr38 = pr.db.ucsc.genes("hg38")
c19 = gr19.coverage(rpm=True, strand=True)
c38 = gr38.coverage(rpm=True, strand=True)
gr38
# +----------------------|-----------|-----------|----------|-----------|----------------|--
# | Chromosome           | Start     | End       | Strand   | Feature   | TranscriptID   | E
# | (int16)              | (int32)   | (int32)   | (int8)   | (int8)    | (int32)        | (
# |----------------------|-----------|-----------|----------|-----------|----------------|--
# | chr1                 | 11873     | 12227     | +        | exon      | NR_046018      | 1
# | chr1                 | 11873     | 12227     | +        | exon      | NR_046018      | 2
# | chr1                 | 11873     | 12227     | +        | exon      | NR_046018      | 3
# | ...                  | ...       | ...       | ...      | ...       | ...            | .
```

```
# | chrY_KZ208924v1_fix | 54153099  | 54153293  | -        | exon      | NR_001537      | 4
# | chrY_KZ208924v1_fix | 54153099  | 54153293  | -        | exon      | NR_001537      | 5
# | chrY_KZ208924v1_fix | 54153099  | 54153293  | -        | exon      | NR_001537      | 6
# +---------------------|-----------|-----------|----------|-----------|----------------|---
# PyRanges object has 863085 sequences from 367 chromosomes.
c38
# chr1 +
# +--------|---------|-------------------|-------------------|--------|-------------------|
# | Runs   | 11873   | 354               | 2182              | 2197   | 159               |
# |--------|---------|-------------------|-------------------|--------|-------------------|
# | Values | 0.0     | 4.634537733826912 | 1.158634433456728 | 0.0    | 1.158634433456728 |
# +--------|---------|-------------------|-------------------|--------|-------------------|
# Rle of length 248919146 containing 7995 elements
# ...
# chrY_KZ208924v1_fix -
# +--------|--------|-------------------|------------|-------------------+
# | Runs   | 4658   | 11067             | 54137374   | 194               |
# |--------|--------|-------------------|------------|-------------------|
# | Values | 0.0    | 1.158634433456728 | 0.0        | 6.951806600740368 |
# +--------|--------|-------------------|------------|-------------------+
# Rle of length 54153293 containing 4 elements
# PyRles object with 643 chromosomes/strand pairs.
c19["chr1"] - c38["chr1"]
# chr1 +
# --
# +--------|---------|--------------------|--------------------|--------------------|-------
# | Runs   | 11868   | 5                  | 354                | 385                | 109
# |--------|---------|--------------------|--------------------|--------------------|-------
# | Values | 0.0     | 4.9974575434747575 | 1.6122841955165352 | 1.3400943382806507 | 5.088
# +--------|---------|--------------------|--------------------|--------------------|-------
# Rle of length 249213345 containing 16054 elements
#
# chr1 -
# --
# +--------|---------|--------------------|-------|---------------------|-------|---------|
# | Runs   | 11873   | 354                | 385   | 109                 | 499   |  ...    |
# |--------|---------|--------------------|-------|---------------------|-------|---------|
# | Values | 0.0     | 1.2493643858686894 | 0.0   | -11.495614382155319 | 0.0   | ...     |
# +--------|---------|--------------------|-------|---------------------|-------|---------|
# Rle of length 249153315 containing 15330 elements
# PyRles object with 2 chromosomes/strand pairs.
```

Run length encoding and arithmetic is a general problem which can be found in many areas of science. Our implementation is the first and only one for Python and we therefore expect it to be much used.

#### Testing PyRanges

To ensure the correctness of PyRanges, we have used the hypothesis library for property based testing (PBT). PBT is a testing method that consists of describing how the data should look and having the computer generate instances based on these blueprints. In our case, the data should look like genomic intervals, which meant our specification was to create a dataframe with columns for chromosomes, starts, ends, and strands, where the chromosomes are drawn from chr1, chr2, . . . , chrM, starts are integers from 0 to maxint32, ends are an integer bigger than start, and strands are from the set ("+", "-").

These instances were used in tests to ensure that the library works as expected. In the general case, one cannot use property based testing to ensure that the library gives the correct result, only that it satisfies some property or invariant and does not throw an error. This is because the solution is not always known. But in our case, we had an oracle to use, namely BEDTools for pyranges and S4Vectors for pyrle.

Using this method of testing actually allowed us to discover unknown bugs in bedtools: https://github.com/arq5x/bedtools2/issues/711, https://github.com/arq5x/bedtools2/issues/710, https://github.com/arq5x/bedtools2/issues/645,

#### Continuous integration

To make pyranges easy to extend, maintain, and contribute to we have used the tool codacy (www.codacy.com) to set up several automatic code tests and reviews for the PyRanges code repository on github. These tests are run each time new code is contributed to the repository, which allows us to see whether a change introduced errors or made the code worse in some way.

Currently, 92% of our code is tested,all tests are passing, and the code quality is A, according to codacy. Furthermore, the wily tool (wily.readthedocs.io) which computes a maintainability index, ranks all our source files as A.

#### Summary

Pyranges is fast, has low memory-requirements, is well-tested and well-written, user-friendly, and the first library of its kind for Python.
