## Supplementaries: time and memory use for "PyRanges: efficient comparison of genomic intervals in Python"

### PyRanges paper supplementaries

This document shows the time and memory usage for all non-basic functions in the ecosystem of PyRanges libraries (PyRanges, PyRles and NCLS). They are compared against their equivalents in Python and R, respectively. The basic PyRanges functionality is compared against R Bioconductor's GenomicRanges and pybedtools. The PyRles-functionality is compared against R Bioconductor's S4Vectors. The NCLS is compared against the intervaltree in the Python bx-python library. For each function the equivalent code from each library is shown.

#### unary

PyRanges functionality that operates on a single PyRanges object. These include functions to sort, cluster and convert ranges into run length encodings (RLE.)

##### Sort

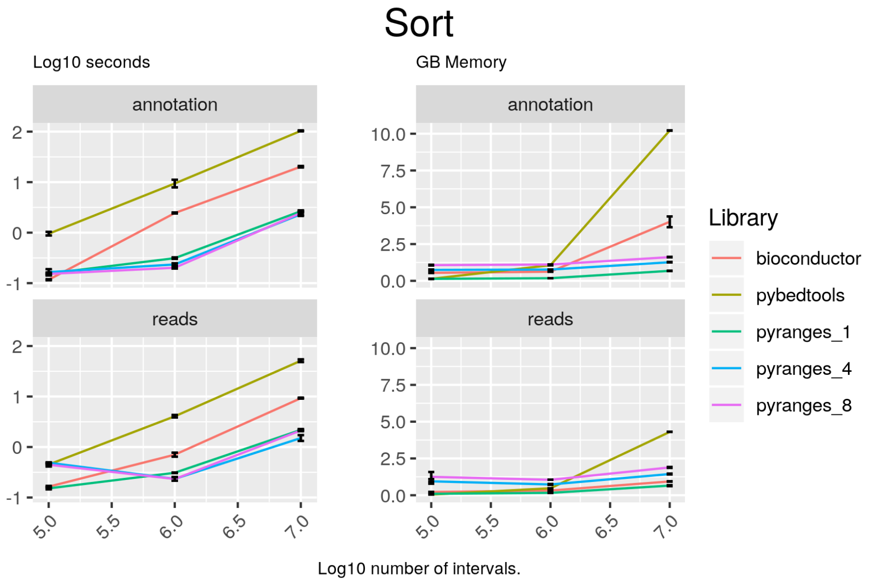

Figure 1: Sort the intervals on Start and End. Comparison of the running time and memory usage for PyRanges versus the equivalent libraries in R and/or Python. In the top row the results for GTF data, while the bottom row shows the results for BED data. The left column shows the time usage, while the right column shows the memory usage. Time is measured in log10 seconds, while memory is measured in GigaBytes (GB).

##### Code

###### pyranges

```
result = gr.sort()
```

###### bioconductor

```
result = sortSeqlevels(gr)
result = sort(result)
```

#### pybedtools

```
result = pb1.sort()
```

##### Cluster

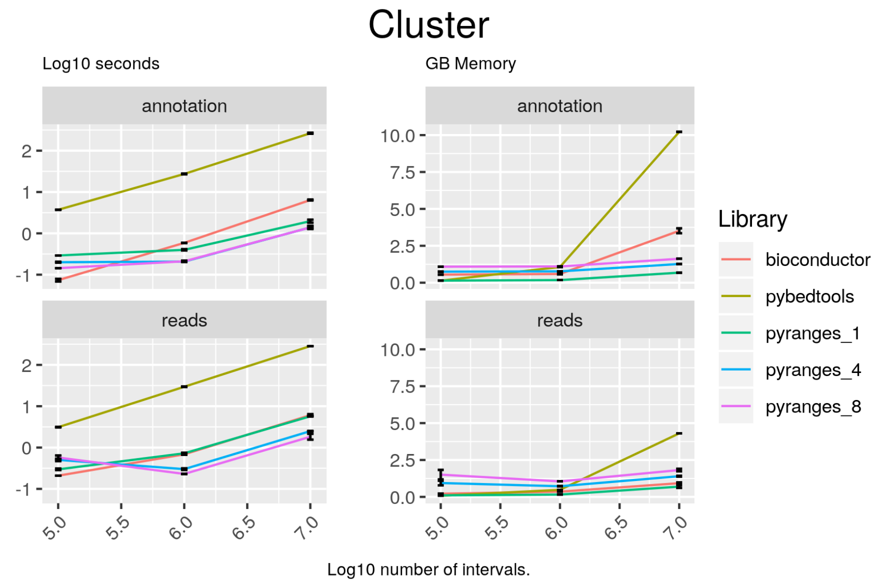

Figure 2: Order intervals by position and merge those overlapping. Comparison of the running time and memory usage for PyRanges versus the equivalent libraries in R and/or Python. In the top row the results for GTF data, while the bottom row shows the results for BED data. The left column shows the time usage, while the right column shows the memory usage. Time is measured in log10 seconds, while memory is measured in GigaBytes (GB).

##### Code

###### pyranges

```
result = gr.cluster(strand="same")
```

###### bioconductor

```
result = reduce(gr)
```

###### pybedtools

```
if extension == "gtf":
    cols_to_keep = [4, 5, 7]
```

```
elif extension == "bed":
    cols_to_keep = [4, 5, 6]

plus = pb1.sort().merge(S="+", c=cols_to_keep, o="first")
minus = pb1.sort().merge(S="-", c=cols_to_keep, o="first")
result = plus.cat(minus, s=True, c=[4, 5, 6], o="first")
```

##### Genomicrange_to_coverage

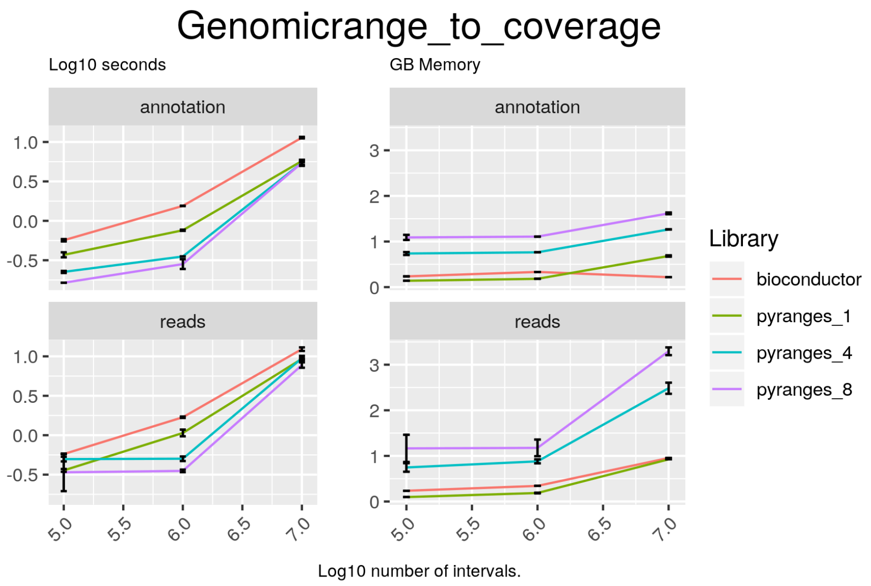

Figure 3: Turn ranges into run length encodings. Comparison of the running time and memory usage for PyRanges versus the equivalent libraries in R and/or Python. In the top row the results for GTF data, while the bottom row shows the results for BED data. The left column shows the time usage, while the right column shows the memory usage. Time is measured in log10 seconds, while memory is measured in GigaBytes (GB).

##### Code

###### pyranges

```
result = gr.coverage(strand="same")
```

###### bioconductor

```
plus = coverage(gr[gr@strand == "+"])
minus = coverage(gr[gr@strand == "-"])
result = c(plus, minus)
```

#### binary

PyRanges functionality that operates on pairs of PyRanges. These functions include functions to find the nearest intervals, find the intersecting intervals, join granges on overlap, set intersect/union and subtract one PyRanges object from another.

##### Overlap

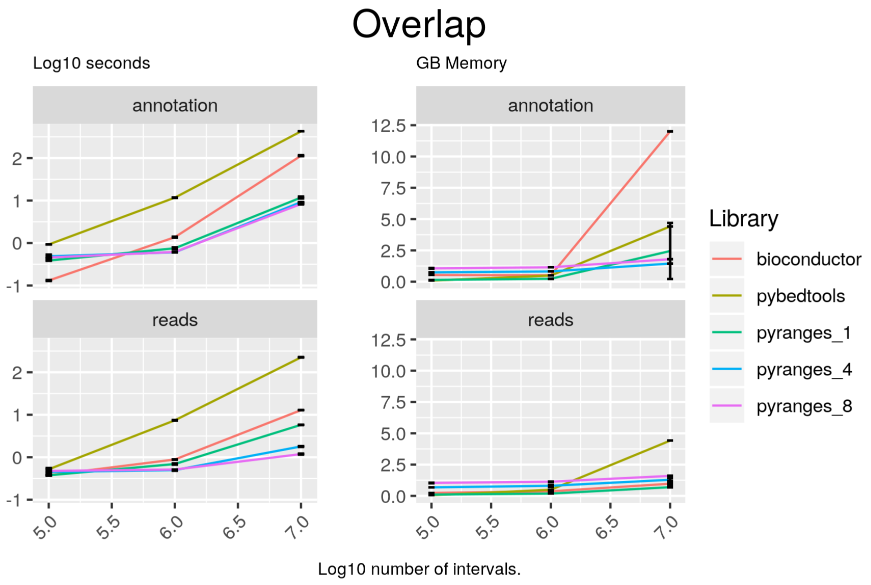

Figure 4: Find the intervals in A overlapping at least one of the intervals in B. Comparison of the running time and memory usage for PyRanges versus the equivalent libraries in R and/or Python. In the top row the results for GTF data, while the bottom row shows the results for BED data. The left column shows the time usage, while the right column shows the memory usage. Time is measured in log10 seconds, while memory is measured in GigaBytes (GB).

##### Code

###### pyranges

```
result = gr2.overlap(gr, strandedness="same")
```

##### bioconductor

```
result = findOverlapPairs(gr2, gr1, ignore.strand = FALSE)
result = first(result)
```

##### pybedtools

```
result = pb2.intersect(pb1, s=True, wa=True)
```

##### Intersect

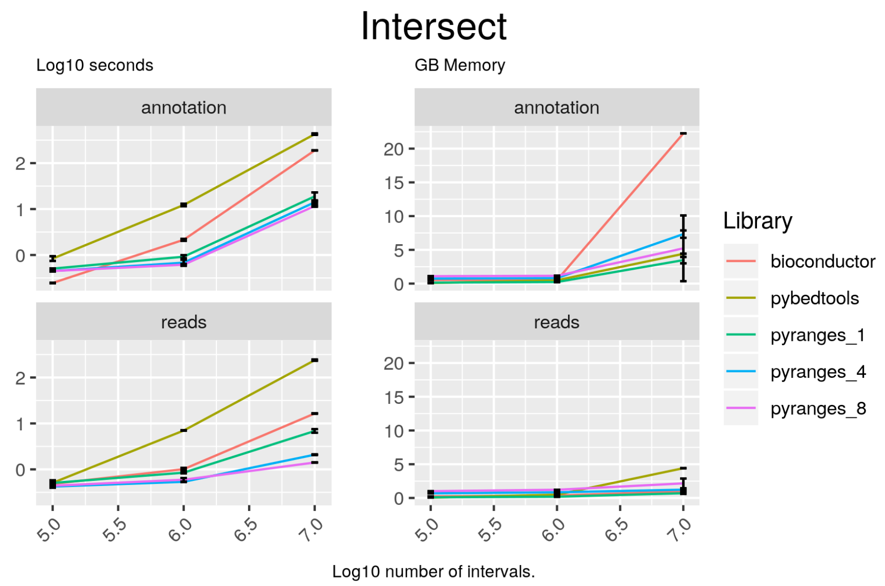

Figure 5: Find overlapping intervals in both datasets. Comparison of the running time and memory usage for PyRanges versus the equivalent libraries in R and/or Python. In the top row the results for GTF data, while the bottom row shows the results for BED data. The left column shows the time usage, while the right column shows the memory usage. Time is measured in log10 seconds, while memory is measured in GigaBytes (GB).

##### Code

###### pyranges

```
result = gr2.intersect(gr, strandedness="same")
```

###### bioconductor

```
pairs = findOverlapPairs(gr2, gr1, ignore.strand = FALSE)
result = pintersect(pairs, ignore.strand = FALSE)
result = result[mcols(result)$hit == TRUE]
```

###### pybedtools

```
result = pb2.intersect(pb1, s=True)
```

##### Nearest

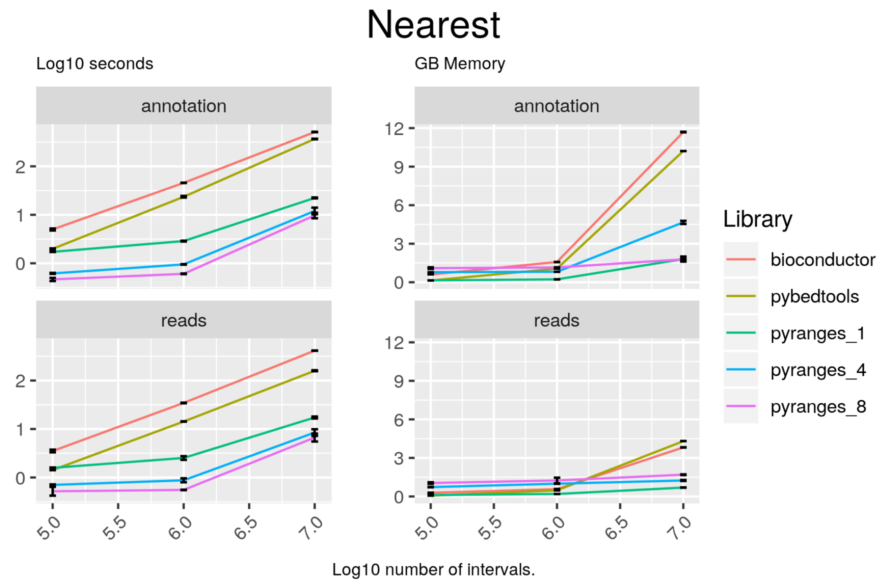

Figure 6: Find the intervals in B closest to those in A. Comparison of the running time and memory usage for PyRanges versus the equivalent libraries in R and/or Python. In the top row the results for GTF data, while the bottom row shows the results for BED data. The left column shows the time usage, while the right column shows the memory usage. Time is measured in log10 seconds, while memory is measured in GigaBytes (GB).

#### Code

##### pyranges

```
result = gr.nearest(gr2, strandedness="same")
```

##### bioconductor

```
result = distanceToNearest(gr2, gr1, ignore.strand = FALSE, select="
    arbitrary")
subject = as.data.frame(gr1[subjectHits(result)])
colnames(subject) = paste0(colnames(subject), "_b")
query = as.data.frame(gr2[queryHits(result)])
df = merge(subject, query, by=0)
df = df[, -1]
df = merge(df, mcols(result)$distance, by=0)
```

```
df = df[, -1]
result = makeGRangesFromDataFrame(df, keep.extra.columns=TRUE)
```

##### pybedtools

```
result = pb2.sort().closest(pb1.sort(), s=True, t="first", d=True)
```

##### Nearest_nonoverlapping

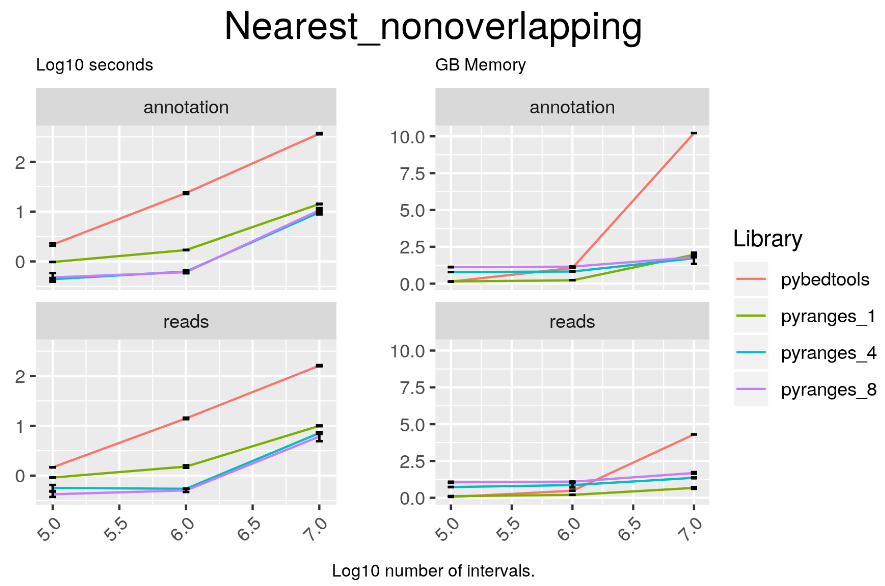

Figure 7: Find the non-overlapping intervals in B closest to those in A. Comparison of the running time and memory usage for PyRanges versus the equivalent libraries in R and/or Python. In the top row the results for GTF data, while the bottom row shows the results for BED data. The left column shows the time usage, while the right column shows the memory usage. Time is measured in log10 seconds, while memory is measured in GigaBytes (GB).

##### Code

##### pyranges

```
result = gr.nearest(gr2, strandedness="same", overlap=False)
```

##### pybedtools

```
result = pb2.sort().closest(pb1.sort(), s=True, t="first", io=True, d=
    True)
```

##### Subtract

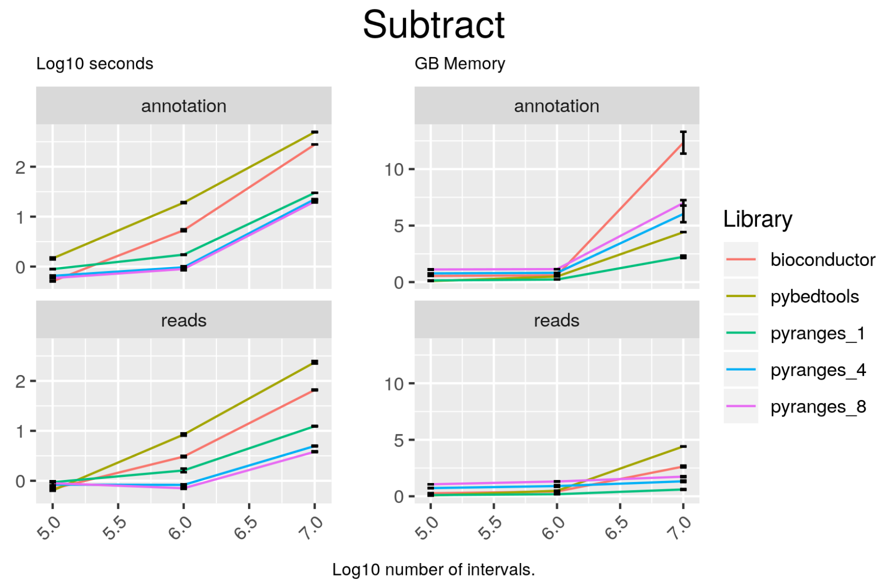

Figure 8: Remove all intervals in B from those in A. Comparison of the running time and memory usage for PyRanges versus the equivalent libraries in R and/or Python. In the top row the results for GTF data, while the bottom row shows the results for BED data. The left column shows the time usage, while the right column shows the memory usage. Time is measured in log10 seconds, while memory is measured in GigaBytes (GB).

##### Code

###### pyranges

```
result = gr2.subtract(gr, strandedness="same")
```

###### bioconductor

```
hits <- findOverlaps(gr2, gr1, ignore.strand = FALSE)
toSubtract <- reduce(extractList(gr1, as(hits, "List")),
    ignore.strand = FALSE)
ans <- unlist(psetdiff(gr2, toSubtract, ignore.strand = FALSE))
result <- subset(ans, width(ans) > 0L)
```

**pybedtools**

```
result = pb2.subtract(pb1, s=True)
```

##### Set_intersect

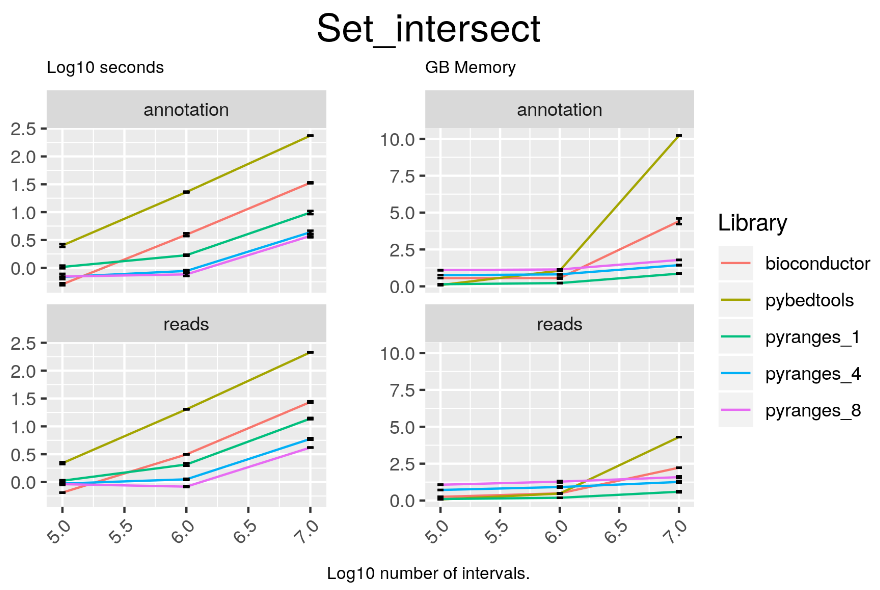

Figure 9: Intersect the set union of the ranges. Comparison of the running time and memory usage for PyRanges versus the equivalent libraries in R and/or Python. In the top row the results for GTF data, while the bottom row shows the results for BED data. The left column shows the time usage, while the right column shows the memory usage. Time is measured in log10 seconds, while memory is measured in GigaBytes (GB).

##### Code

##### pyranges

```
result = gr2.set_intersect(gr, strandedness="same")
```

##### bioconductor

```
result = intersect(gr2, gr1)
```

##### pybedtools

```
sc = pb1.sort().merge(s=True, c=[4, 5, 6], o="first")
```

```
if extension == "gtf":
    cols_to_keep = [4, 5, 7]
elif extension == "bed":
    cols_to_keep = [4, 5, 6]

sb = pb2.sort().merge(s=True, c=cols_to_keep, o="first")
result = sc.intersect(sb, s=True)
```

##### Set_union

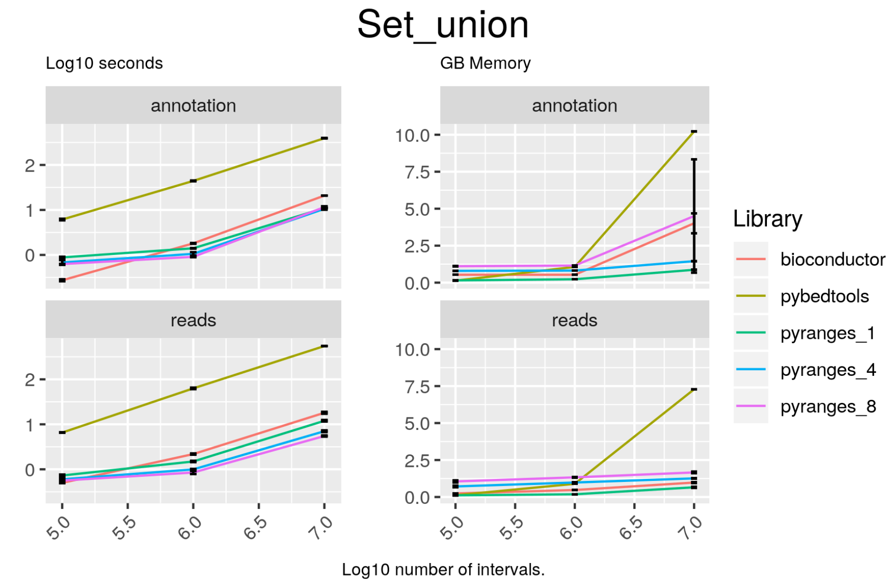

Figure 10: Concatenate the datasets and cluster them afterwards. Comparison of the running time and memory usage for PyRanges versus the equivalent libraries in R and/or Python. In the top row the results for GTF data, while the bottom row shows the results for BED data. The left column shows the time usage, while the right column shows the memory usage. Time is measured in log10 seconds, while memory is measured in GigaBytes (GB).

##### Code

##### pyranges

```
result = gr2.set_union(gr, strandedness="same")
```

##### bioconductor

```
result = union(gr1, gr2)
```

##### pybedtools

```
sc = pb1.sort().merge(s=True, c=[4, 5, 6], o="first")
```

```
if extension == "gtf":
    cols_to_keep = [4, 5, 7]
elif extension == "bed":
    cols_to_keep = [4, 5, 6]

sb = pb2.sort().merge(s=True, c=cols_to_keep, o="first")
catted = sc.cat(sb, s=True, c=[4, 5, 6], o="first").sort()
result = catted.merge(s=True, c=[4, 5, 6], o="first")
```

##### Jaccard

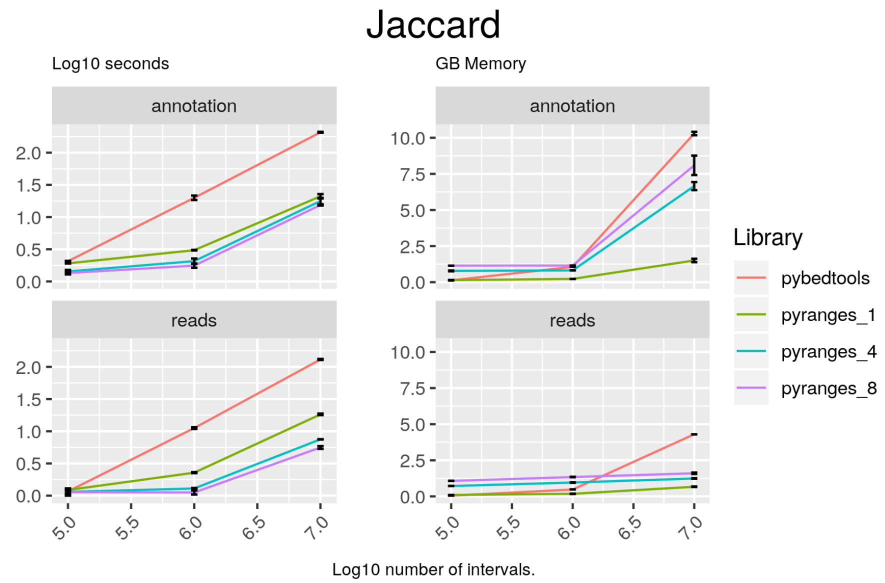

Figure 11: Find similarity between sets based on intersections. Comparison of the running time and memory usage for PyRanges versus the equivalent libraries in R and/or Python. In the top row the results for GTF data, while the bottom row shows the results for BED data. The left column shows the time usage, while the right column shows the memory usage. Time is measured in log10 seconds, while memory is measured in GigaBytes (GB).

##### Code

###### pyranges

```
result = gr2.stats.jaccard(gr)
```

###### pybedtools

```
result = pb2.sort().jaccard(pb1.sort())
```

##### Join

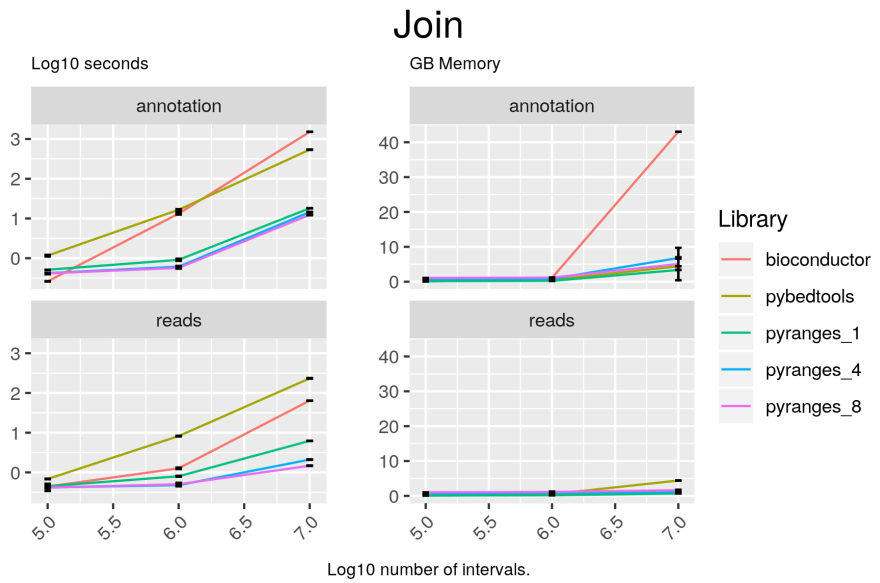

Figure 12: Find the overlapping intervals and combine their data. Comparison of the running time and memory usage for PyRanges versus the equivalent libraries in R and/or Python. In the top row the results for GTF data, while the bottom row shows the results for BED data. The left column shows the time usage, while the right column shows the memory usage. Time is measured in log10 seconds, while memory is measured in GigaBytes (GB).

##### Code

###### pyranges

```
result = gr2.join(gr, strandedness="same")
```

###### pybedtools

```
result = pb2.intersect(pb1, wao=True, s=True)
```

###### bioconductor

```
result <- findOverlapPairs(gr1, gr2, ignore.strand = FALSE)
df1 = as.data.frame(first(result))
```

```
df2 = as.data.frame(second(result))
colnames(df2) = paste0(colnames(df2), "_b")
df = merge(df1, df2, by=0)
result = makeGRangesFromDataFrame(df, keep.extra.columns=TRUE)
```

#### rle

Arithmetic operations on RLEs. These include add, subtract, divide and multiply.

##### Rle_divide

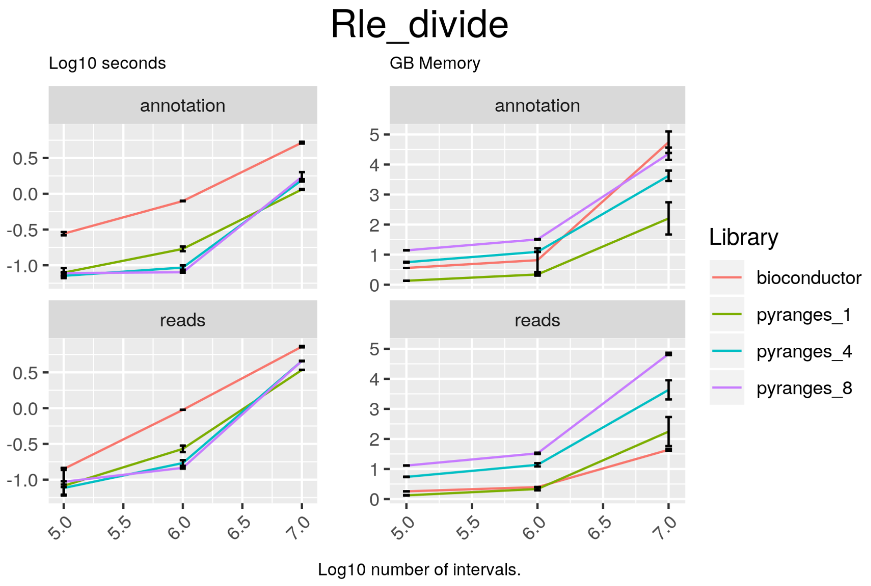

Figure 13: Divide one Rle object by another. Comparison of the running time and memory usage for PyRanges versus the equivalent libraries in R and/or Python. In the top row the results for GTF data, while the bottom row shows the results for BED data. The left column shows the time usage, while the right column shows the memory usage. Time is measured in log10 seconds, while memory is measured in GigaBytes (GB).

##### Code

##### bioconductor

```
result = c(c1p / c2p, c1m / c2m)
```

##### pyranges

```
result = c1 / c2
```

##### Rle_add

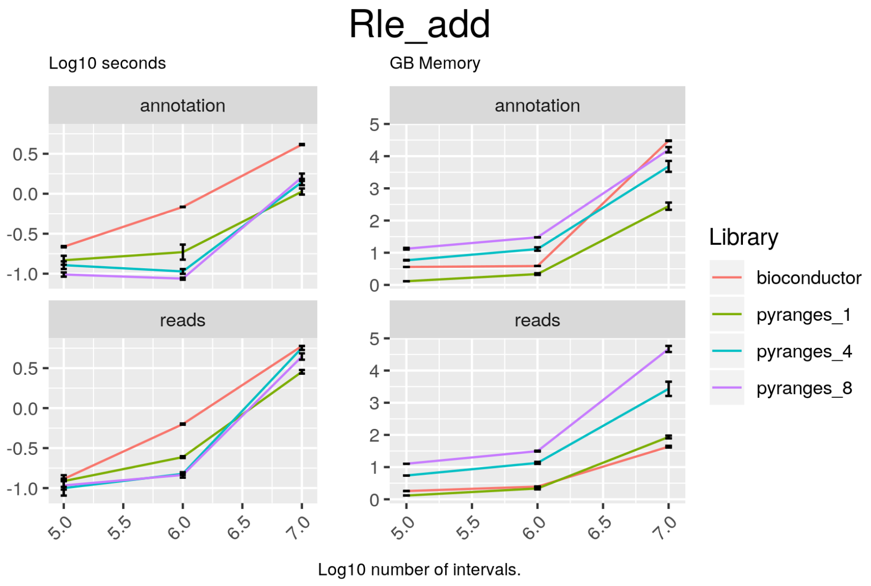

Figure 14: Add two Rle objects. Comparison of the running time and memory usage for PyRanges versus the equivalent libraries in R and/or Python. In the top row the results for GTF data, while the bottom row shows the results for BED data. The left column shows the time usage, while the right column shows the memory usage. Time is measured in log10 seconds, while memory is measured in GigaBytes (GB).

##### Code

###### bioconductor

```
result = c(c1p + c2p, c1m + c2m)
```

###### pyranges

```
result = c1 + c2
```

##### Rle_subtract

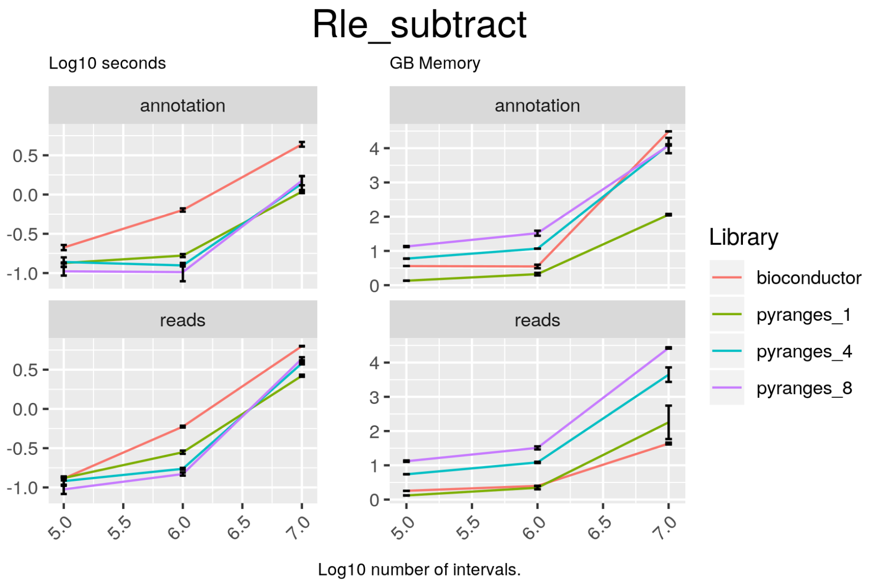

Figure 15: Subtract one Rle object from another. Comparison of the running time and memory usage for PyRanges versus the equivalent libraries in R and/or Python. In the top row the results for GTF data, while the bottom row shows the results for BED data. The left column shows the time usage, while the right column shows the memory usage. Time is measured in log10 seconds, while memory is measured in GigaBytes (GB).

#### Code

##### bioconductor

```
result = c(c1p - c2p, c1m - c2m)
```

##### pyranges

```
result = c1 - c2
```

##### Rle_multiply

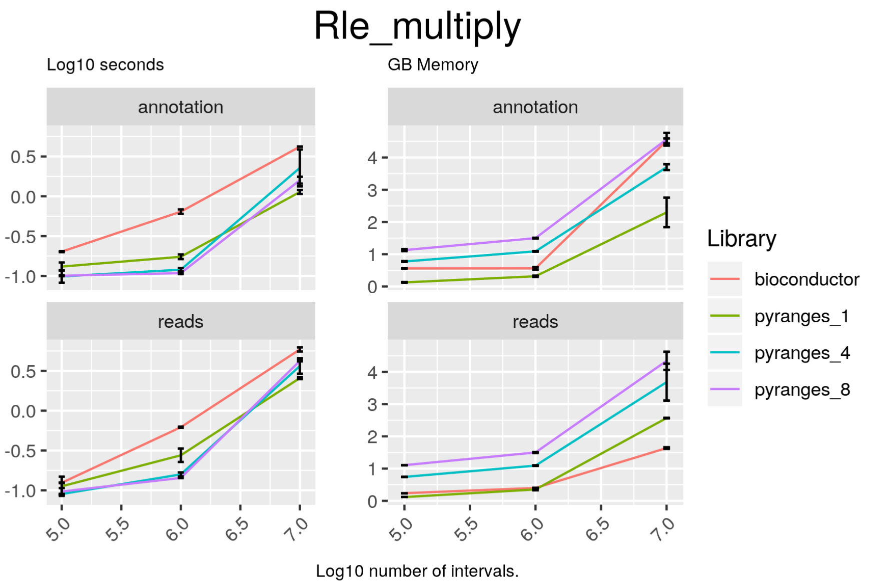

Figure 16: Multiply two Rle objects. Comparison of the running time and memory usage for PyRanges versus the equivalent libraries in R and/or Python. In the top row the results for GTF data, while the bottom row shows the results for BED data. The left column shows the time usage, while the right column shows the memory usage. Time is measured in log10 seconds, while memory is measured in GigaBytes (GB).

##### Code

##### bioconductor

```
result = c(c1p * c2p, c1m * c2m)
```

##### pyranges

```
result = c1 * c2
```

##### tree

Operations for building and finding overlaps using a tree.

###### Tree_build

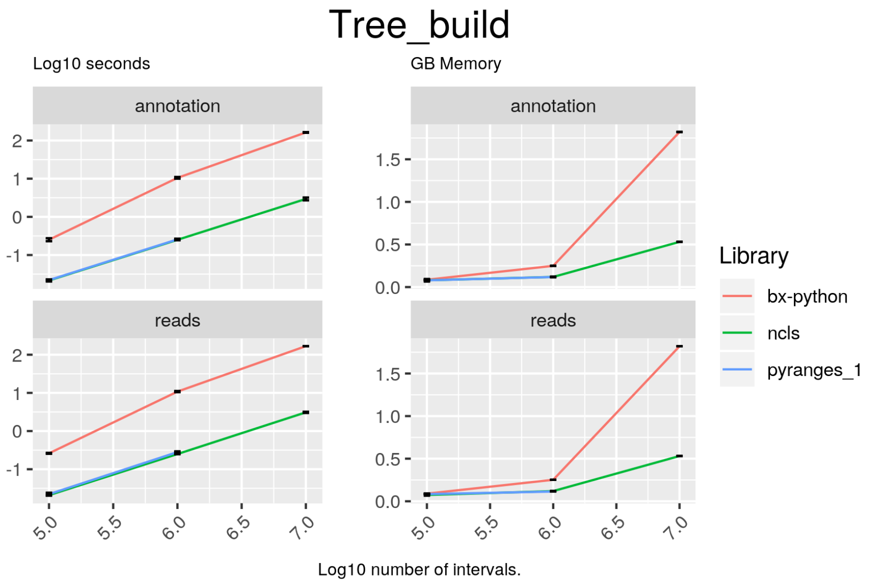

Figure 17: Create a tree from a collection of intervals. Comparison of the running time and memory usage for PyRanges versus the equivalent libraries in R and/or Python. In the top row the results for GTF data, while the bottom row shows the results for BED data. The left column shows the time usage, while the right column shows the memory usage. Time is measured in log10 seconds, while memory is measured in GigaBytes (GB).

###### Code

###### ncls

```
tree = NCLS(df2.Start.values, df2.End.values, df2.index.values)
```

###### bx-python

```
tree = IntervalTree()
for start_, end_ in zip(df2.Start, df2.End):
    tree.add(start_, end_)
```

##### Tree_overlap

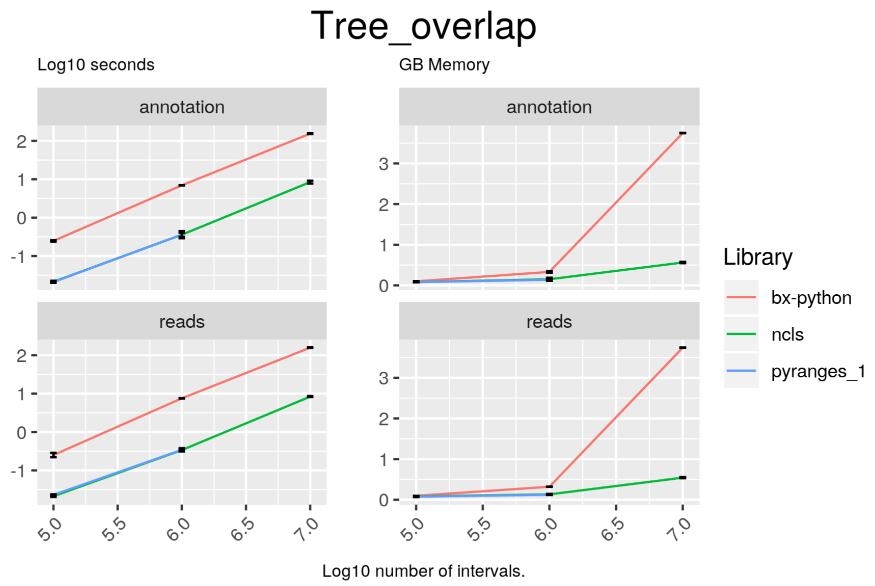

Figure 18: Search in a tree using a collection of intervals. Comparison of the running time and memory usage for PyRanges versus the equivalent libraries in R and/or Python. In the top row the results for GTF data, while the bottom row shows the results for BED data. The left column shows the time usage, while the right column shows the memory usage. Time is measured in log10 seconds, while memory is measured in GigaBytes (GB).

#### Code

##### ncls

```
result = tree.all_overlaps_self(df1.Start.values, df1.End.values, df1.
    index.values)
result = df2.iloc[result]
```

##### bx-python

```
result = []
for start_, end_ in zip(df1.Start, df1.End):
    result.append(tree.search(start_, end_))
```

## io

Functions to read files into PyRanges.

##### Read_bed

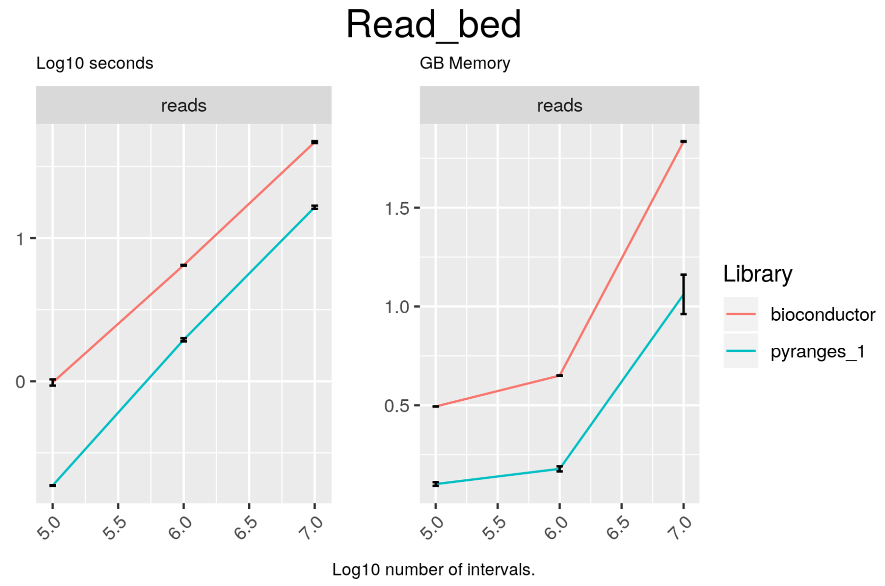

Figure 19: Read a bed file into a GenomicRanges object. Comparison of the running time and memory usage for PyRanges versus the equivalent libraries in R and/or Python. In the top row the results for GTF data, while the bottom row shows the results for BED data. The left column shows the time usage, while the right column shows the memory usage. Time is measured in log10 seconds, while memory is measured in GigaBytes (GB).

##### Code

###### pyranges

```
result = pr.read_bed(f)
```

###### bioconductor

```
result = import(file)
```

##### Read_bam

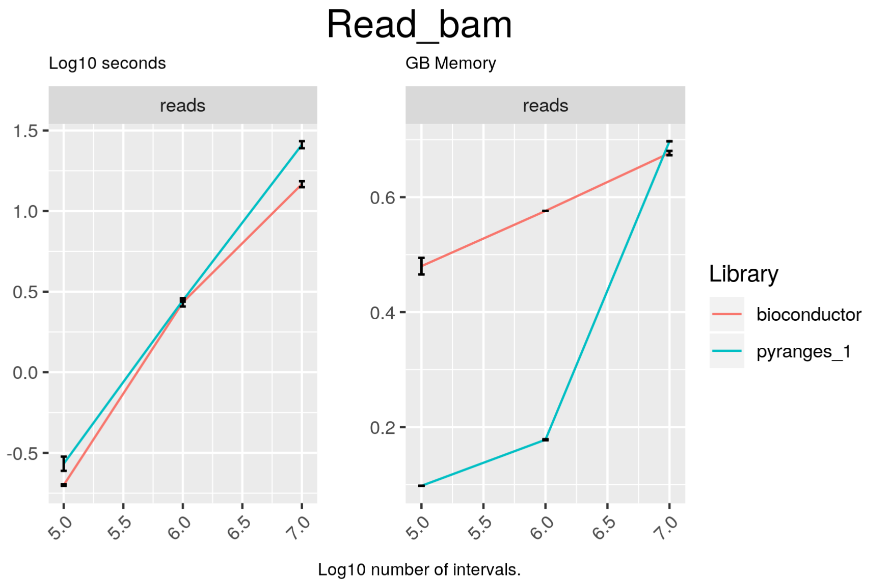

Figure 20: Read a bam file into a GenomicRanges object. Comparison of the running time and memory usage for PyRanges versus the equivalent libraries in R and/or Python. In the top row the results for GTF data, while the bottom row shows the results for BED data. The left column shows the time usage, while the right column shows the memory usage. Time is measured in log10 seconds, while memory is measured in GigaBytes (GB).

##### Code

##### pyranges

```
result = pr.read_bam(f)
```

##### bioconductor

```
result = import(file)
```

#### Read_gtf

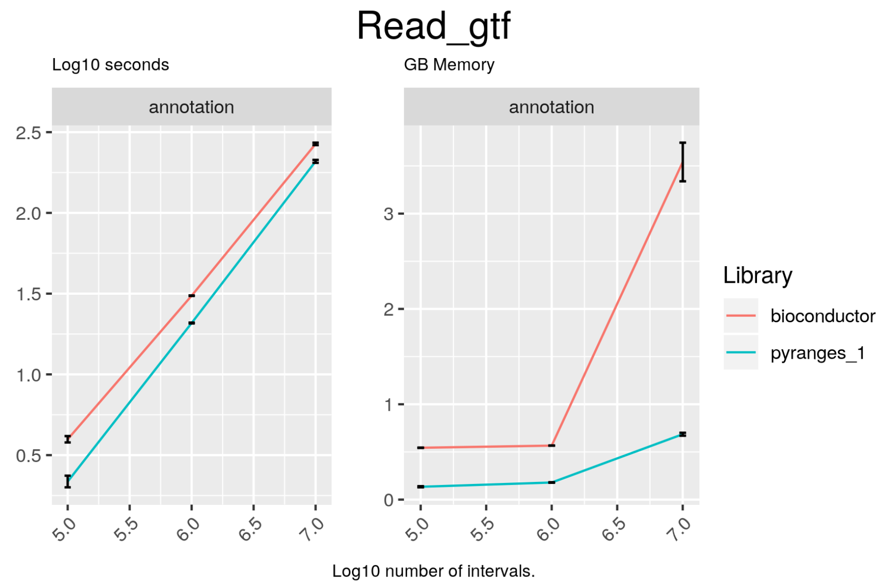

Figure 21: Read a gtf file into a GenomicRanges object. Comparison of the running time and memory usage for PyRanges versus the equivalent libraries in R and/or Python. In the top row the results for GTF data, while the bottom row shows the results for BED data. The left column shows the time usage, while the right column shows the memory usage. Time is measured in log10 seconds, while memory is measured in GigaBytes (GB).

#### Code

##### pyranges

```
result = pr.read_gtf(f, annotation="ensembl")
```

##### bioconductor

```
result = import(file)
```
